# Combinatorial bnAb and T Cell Immunization Enhances Post-Infection Control of SHIV_SF162p3_ Viremia in Rhesus Macaques

**DOI:** 10.64898/2026.11.04.764342

**Authors:** Diane G. Carnathan, Chiamaka A. Enemuo, Julia T. Ngo, Luis E. Jimenez, Yury Choe, Jennifer B. Silverman, Stanca Ciupe, Guido Silvestri, Sudhir P. Kasturi, Robert C. Gallo, George K. Lewis

**Author notes:** Corresponding authors George K. Lewis, Ph.D., Institute of Human Virology, 725 West Lombard Street, Baltimore, Maryland, 21201. Robert C. Gallo, M.D., Institute of Human Virology, 725 West Lombard Street, Baltimore, Maryland, 21201.^6^,^4^. Robert C. Gallo, M.D., ^1^Institute for Translational Virology and Innovation, Department of Internal Medicine, Morsani College of Medicine, University of South Florida, Tampa, FL, USA, and Tampa General Hospital Cancer Institute, Tampa, FL, USA. ^1^,^1^,^1^,^1^,^1^,^1^,^4^,^2^,^1^. **Author Contributions:** Designed Research (DGC, GS, RCG, SPK, GKL). Performed Research (DCG, CAE, JTN, LEJ, YC, JBS, SPK, GKL). Analyzed Data (DCG, GS, SPK, SC, GKL). Wrote the paper (DCG, GS, GKL). **Competing Interests** The authors declare no competing interests.

## Abstract

An ideal HIV-1 vaccine should provide broad protection against most HIV-1 variants, control early dissemination of infected cells after virus acquisition, and avoid the risk that vaccine-induced, HIV-specific CD4+ T cells serve as targets for viral transmission. In this study, we modeled this complex host-virus interaction in rhesus macaques challenged with a single high dose of SHIV_SF162p3_. Specifically, we evaluated the impact of T cell immunization with a human adenovirus-5-based candidate vaccine encoding an SIV Gag-Tat fusion (AdHu5_SIVGag/Tat_) alongside a partially protective, limiting dose of the broadly neutralizing antibody (bnAb) PGT121. Immunization with AdHu5_SIVGag/Tat_ elicited Gag-specific CD4+ and CD8+ T cell responses in PBMC and mucosal tissue. T cell vaccination did not affect PGT121-mediated protection against SHIV_SF162p3_ acquisition. However, independently of vaccination, constitutively activated CD4+α4β7+Ki-67+ T cells in rectal tissue attenuated PGT121-mediated protection. Crucially, a unique pattern of post-infection control of SHIVSF162p3 viremia emerged in AdHu5_SIVGag/Tat_-immunized, PGT121-treated animals, characterized by an increased eclipse phase, lower peak viremia, and a late-phase decay in steady-state viremia. These results suggest potential synergy between bnAbs and AdHu5_SIVGag/Tat_-induced antiviral T cell responses, with significant implications for HIV-1 vaccine development and early interventions following infection.

## Introduction

After almost forty years of intense effort, a safe and effective HIV-1 vaccine remains elusive. Driven by extensive preclinical research, nine HIV-1 vaccine efficacy trials have been completed since 1999, with only one trial, RV144, [2] reporting weak efficacy (reviewed in [3]). In addition, increased HIV-1 transmission was observed in two trials, HVTN 502 [4] and HVTN 503 [5], further complicating HIV-1 vaccine development. Despite the lack of significant, durable protection from the vaccines used in these nine efficacy trials, subgroup analyses have provided important hypothesis-generating clues about vaccine-elicited protection that can be evaluated in nonhuman primate (NHP) models. While this approach is not optimal, as shown by the modest ability of RM studies to predict outcomes in human efficacy trials (reviewed in [6]), it remains the only way to evaluate HIV-1 vaccine concepts outside clinical trials. Despite this caveat, five consistent themes emerge from vaccine, passive immunization, and natural history studies in NHPs and humans.

First, antibodies can prevent viral transmission if they effectively neutralize founder viruses and are present at protective levels during exposure. This has been consistently demonstrated in NHP studies (reviewed in [7, 8]) and confirmed in two Phase 2b efficacy trials of the bnAb VRC01 to prevent HIV-1 acquisition in healthy volunteers [9]. Although these trials showed no overall efficacy, subgroup analyses revealed that VRC01 neutralized both transmitting and non-transmitting HIV-1 variants, with non-transmitting variants neutralized at IC_80_ ≤ 1ug/ml and transmitting variants at IC_80_ ≤ 3ug/ml [10, 11]. These findings illustrate the pharmacologically relevant bnAb levels required for protection in both active and passive immunization, although viral diversity remains a major challenge. Despite these issues, virus neutralization is the only known immune mechanism that reliably blocks HIV/SIV/SHIV acquisition at exposure, and it is widely accepted that an effective HIV-1 vaccine should induce protective bnAb levels.

Second, bnAb levels in RM at the time of SHIV/SIV challenge that incompletely protect against infection result in post-infection control of viremia, characterized by an increased “eclipse” phase (time to first detectable viremia), decreased peak viremia, and increased setpoint control of viremia (16-24). While non-neutralizing antibodies (nnAbs) also enable post-infection control of viremia in NHP passive immunization models, they do not block high-dose SHIV challenges (Field (6, 24-30)). This result is consistent with many NHP studies over the years, in which vaccine-elicited nnAbs correlate with increased resistance to low-dose SHIV/SIV challenges (Field (31-37)). Nevertheless, only bnAbs exhibit the breadth and potency to reliably prevent infection at the time of virus exposure, regardless of inoculum size. These observations provide additional support for bnAb-based approaches to block HIV-1 acquisition and point to the ability of sub-blocking concentrations of bnAbs to elicit post-infection control of viremia, long after bnAb concentrations have decayed to background levels.

Third, passive immunization of SIV/SHIV-infected RM with bnAbs shortly after infection can lead to long-term viral control. Early studies showed that polyclonal immunoglobulin against SIV (SIVIG) altered the disease course in some RM over 67 weeks, accelerating neutralizing antibody production and maintaining Gag-specific CD8+ CTLs for more than a year [12]. In SHIV-infected infants, early bnAb administration resulted in undetectable viral loads at 6 months post-infection, with no rebound or virus-specific T cells at study end, indicating prevention of viral outgrowth and reservoir formation [13]. Similar results were observed with a single bnAb dose post-challenge [14]. More recent studies showed that infusing two bnAbs three days after SHIV challenge, followed by two weekly bnAb infusions, achieved long-term control for up to six years and correlated with virus-specific CD8+ T cells, suggesting a “vaccinal” effect [15, 16]. Thus, early bnAb treatment appears to enhance the ability of CD8+ T cells to achieve durable control without ART, compared with early ART alone [16]. Little is known about how often or the mechanisms that enable bnAbs and virus-specific CD8+ T cells to interact in early intervention models (reviewed in [17, 18]). Very recently, synergy between bnAbs and virus-specific CD8+ T cells for in vitro control of HIV-1 was reported [19], and an in vivo RM study [20] showed that partially protective bnAb levels enhance SIV replication arrest elicited by the well-known RhCMV/SIV vaccine [21–25].

Fourth, studies in humans and RMs demonstrate that CD8+ T cells are crucial for controlling HIV-1/SIV/SHIV after infection, regardless of antibody responses to the envelope glycoprotein. Early research linked viremic control to the emergence of HIV-1-specific CD8+ T cells in humans [26, 27] and RMs [28], a link supported by the rapid development of escape variants in HIV-1-infected individuals [29–31] and SIV-infected RMs [32–34]. Additionally, HIV-positive individuals who spontaneously suppress viral loads below 50 copies/ml without ART, known as elite controllers (EC), further suggest a role for CD8+ T cells in viral regulation. These individuals often have specific Class I MHC alleles [35, 36], and strong evidence indicates that HIV-1-specific CD8+ T cells contribute to viral control in EC populations [37–40], as reviewed in [41]. Moreover, vaccinating RMs with a rhesus macaque CMV vector encoding SIV proteins induces CD8+ effector memory T cells [21] that correlate with effective immune control of early SIVmac239 infections [22–25, 42], with about half achieving viral clearance [23]. Others show long-term control of viremia, akin to that in human EC [23–25, 42]. These findings propose that HIV-1-specific CD8+ T cells can significantly contribute to controlling HIV-1 variants that evade broadly neutralizing antibodies, as also indicated by early bnAb intervention studies [15, 16].

Fifth, several studies suggest that vaccine-elicited CD4+ T cells can increase HIV/SHIV/SIV replication and acquisition, thereby tempering vaccine effectiveness, presumably by increasing the number of CD4+ T cells susceptible to infection (reviewed in [43]). This effect was implicated in the STEP/Phambili HIV-1 vaccine efficacy trials, where immunization with a mixture of three AdHu5 vectors encoding HIV-1 *gag*, *pol*, or *nef* increased acquisition among subsets of men in both studies [44, 45]. In the absence of anti-Env antibodies, vaccine-elicited CD4+ T cells were implicated as a mechanism of increased acquisition ([46–49], summarized in [50]). However, it remains to be seen whether host factors also contribute to increased acquisition [51]. Several RM studies using SIV or SHIV challenges suggest that vaccine-elicited CD4+ T cell responses correlate with increased acquisition [49], decreased vaccine efficacy [52–54], or increased viral replication after infection [55, 56].

These five themes outline an emerging relationship between antibody-mediated protection against acquisition and post-infection control of viremia by CD8+ T cells, supporting a combined bnAb/T cell HIV-1 vaccine strategy. This approach should prevent infection in exposed individuals and reduce viral loads in infected vaccinees to levels below the transmission threshold [57–60]. Ideally, the vaccine will sustain viremia control below the transmission threshold without ART or work synergistically with less ART to prevent transmission. This strategy should also reduce morbidity and mortality among vaccinees with breakthrough infections.

Based on these considerations, we designed an RM study to evaluate a combinatorial bnAb/T cell vaccination strategy in an acute SHIV_SF162p3_ infection model. Because no current vaccines elicit bnAbs, we modeled bnAb-mediated protection by passively immunizing RM with a limiting dose of the bnAb PGT121, which protected approximately two-thirds of RM against SHIV_SF162p3_ infection in this study. We modeled T cell vaccination with a human adenovirus-5 vaccine encoding an SIV Gag-Tat fusion (AdHu5_SIVGag/Tat_), which our team used to show that activated rectal CD4+CCR5+ T cells predicted increased SIV_mac239_ acquisition in RM [56]. We made the following observations.

Passive immunization with PGT121 alone protected 66.7% (6/9) of RM from SHIV_SF162p3_ infection, whereas no protection was observed in RM actively immunized with AdHu5_SIVGag/Tat_ alone. Combined active and passive immunization with AdHu5_SIVGag/Tat_ and PGT121 also conferred 66.7% protection against SHIV_SF162p3_ infection, with no effect, positive or negative, on PGT121’s ability to block infection. Post hoc analyses showed that higher baseline frequencies of constitutive mucosal CD4+α4β7+Ki-67+ T cells correlated with increased susceptibility to infection across animals receiving PGT121, independent of vaccination status. Breakthrough infections in the PGT121-only group displayed a delayed eclipse phase, reduced peak viral loads, and moderate setpoint viral control relative to controls. RM immunized with AdHu5_SIVGag/Tat_ alone showed characteristic post-infection control of viremia, marked by lower peak viremia and reduced steady-state viral loads. By contrast, combined active immunization with AdHu5_SIVGag/Tat_ and passive immunization with PGT121 revealed a unique, synergistic pattern of viral control, characterized by an extended eclipse phase, lower peak viremia, and a statistically significant late-stage decay of viremia, approaching the limit of detection in the final weeks of the study.

Taken together, these results support the concept that combinatorial active and passive immunization with T cell vaccines and bnAbs results in synergistic early control of SHIV_SF162p3_ viremia. Our results complement a recent report using a similar approach in the SIV system, suggesting synergy between T cell vaccination and bnAb prophylaxis for enhanced control of SHIV/SIV early in infection [20]. Results of these studies are discussed in the context of antigen cross-presentation and “vaccinal” effects in the post-infection control of retroviral replication and disease.

## Results

### Study Design

We designed the study shown in Figure 1 to evaluate the impact of T cell vaccination on protection against a single, high-dose rectal challenge with SHIV_SF162p3,_ mediated by a limiting dose of the highly potent bnAb PGT121 expressed in plants [61]. Based on prior PK studies [62] and in-house confirmation, we used a PGT121 dose predicted to protect approximately sixty percent of RM from a single, high-dose rectal challenge with SHIV_SF162p3_. The T cell vaccine was a human adenovirus-5 vector encoding a Gag-Tat fusion protein (AdHu5_SIVGag/Tat_) based on the SIV_mac239_ infectious molecular clone [56]. Previous studies from our team [56] showed that SIVGag/Tat-immunized RM that became infected after repeat low-dose rectal challenges with SIV_mac239_ exhibited lower setpoint control of viremia, although there was no protection against infection. Vaccinated RM that became infected had higher frequencies of pre-challenge rectal CD4+CCR5+DR+ T cells, consistent with earlier studies suggesting a relationship between HIV-1 acquisition and vaccine-specific CD4+ T cells [5, 44–49]. Increased day 7 viremia also correlated directly with higher pre-challenge frequencies of rectal CD4+CCR5+Ki-67+ T cells [56]. Interestingly, despite these effects on infection susceptibility and early viral replication, vaccinated RM exhibited lower setpoint post-infection viremia control than unvaccinated RM [56]. Thus, despite increased infection and early viral loads, the AdHu5_SIVGag/Tat_ vaccine elicited post-infection control of viremia [56]. Based on these considerations, the study scheme shown in Figure 1 was designed to test whether T cell vaccination impacts bnAb efficacy or vice versa.

**Figure 1.**
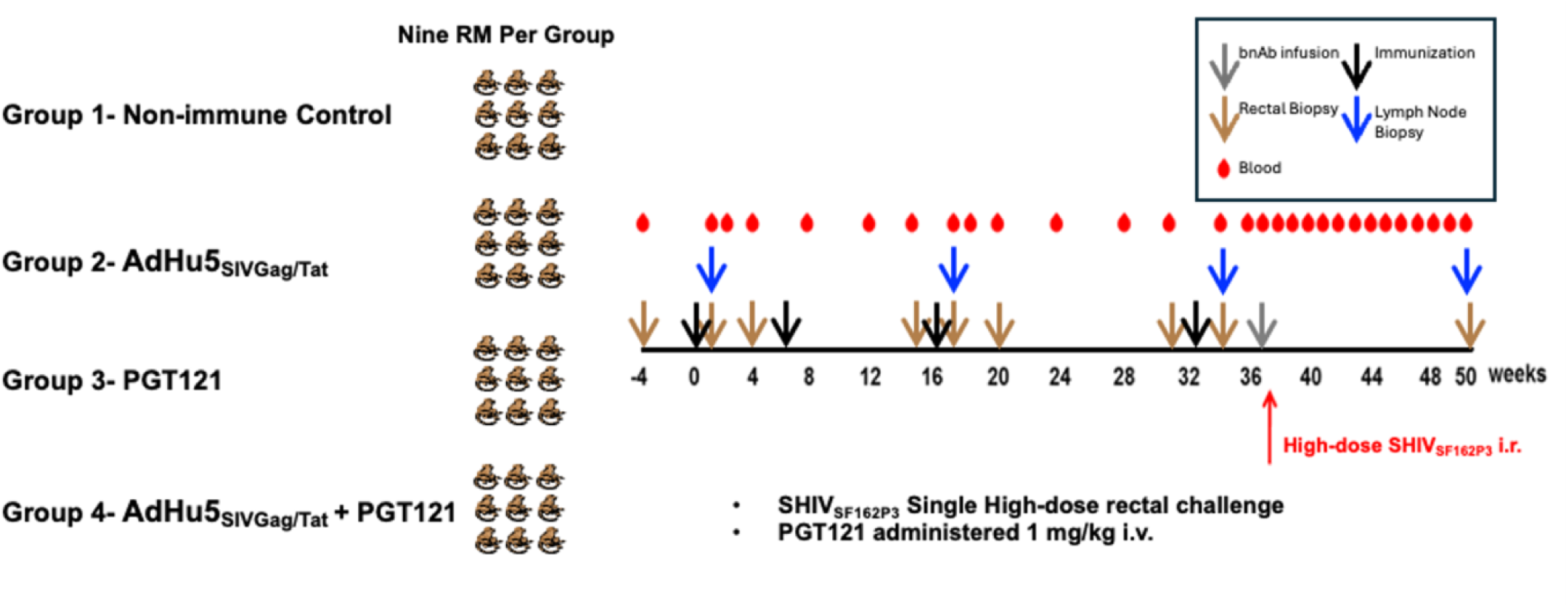
Study design groups and general schedule. Schematic representation of AdHu5 immunization, PGT121 administration, and challenge. 36 Mamu*B08- and *B17-adult Indian RMs were divided into four groups of nine animals. 18 RMs were immunized with AdHu5_SIVGat/Tat_ three times with 16 weeks in between (blue arrows) and 18 animals remained unimmunized. Four weeks following the final immunization, 18 animals (nine AdHu5_SIVGat/Tat_ vaccinated and nine unvaccinated) were administered PGT121 intravenously (gray arrow). The following day all 36 RMs were challenged intrarectally with a single dose of SHIV_162P3_ (red arrow). All animals were followed for up to 14 weeks after the challenge. Blood, LN FNA and RB were collected at intervals throughout the study.

Thirty-six adult Indian RMs, Mamu-B*08 - and B*17-positive, were randomized into four groups of nine RMs per group. Specimen collections and active and passive immunizations were performed according to the schedule shown in Figure 1. Group 1 served as the infection control group. Groups 2 and 4 were immunized with AdHu5_SIVGag/Tat_ three times at sixteen-week intervals (black arrows, Figure 1). Four weeks after the last immunization of Groups 2 and 4, Groups 3 and 4 were passively immunized intravenously with a 1.0 mg/kg dose of PGT121 (gray arrow, Figure 1). All thirty-six RMs were challenged intrarectally the following day with a single, high-dose SHIV_SF162p3_ (red arrow, Figure 1). Plasma viral loads were monitored for 14 weeks post-challenge. Blood (red drops, Figure 1), LN FNA (blue arrows, Figure 1), and rectal biopsies (brown arrows, Figure 1) were also collected at intervals throughout the study as indicated.

### T cell Vaccination Does Not Abrogate or Enhance Protection Against SHIV_SF162p3_ Infection Elicited by Limiting Dose of bnAb PGT121

Figure 2 shows the frequencies of protection against SHIV_SF162p3_ infection among Groups 2, 3, and 4. Eight of nine RM in the challenge control group (Group 1) and nine of nine RM in the AdHu5_SIVGag/Tat_-immunized group (Group 2) became infected, consistent with our prior study showing that AdHu5_SIVGag/Tat_ immunization did not protect against low-dose SIV_mac239_ challenges [56]. By contrast, six of nine RM were protected from infection by receiving either PGT121 (Group 3) or PGT121 plus vaccination with AdHu5_SIVGag/Tat_ (Group 4). Three RM in each of Groups 3 and 4 became infected, and there was no significant difference in acquisition between the two groups (statistical analysis in Table 1). The sole correlate of protection against SHIV_SF162p3_ infection was administration of the bnAb PGT121. Prior immunization with AdHu5_SIVGag/Tat_ did not attenuate the ability of a limiting dose of PGT121 to protect against a single, high-dose rectal challenge with SHIV_SF162p3_, although AdHu5_SIVGag/Tat_ vaccination elicited CD4+ and CD8+ SIV Gag-specific T cell responses in PBMC measured by antigen-specific intracellular cytokine production (Figures S1 and S2). AdHu5_SIVGag/Tat_ vaccination also elicited Gag-CM9 Mamu-A*01 Tetramer-Binding CD8+ T cells in PBMC (Figure S3A), and their mean frequency in rectal biopsies three weeks before SHIV_SF162p3_ challenge was 1.85% of total CD8+ T cells (Figure S3B).

**Figure 2.**
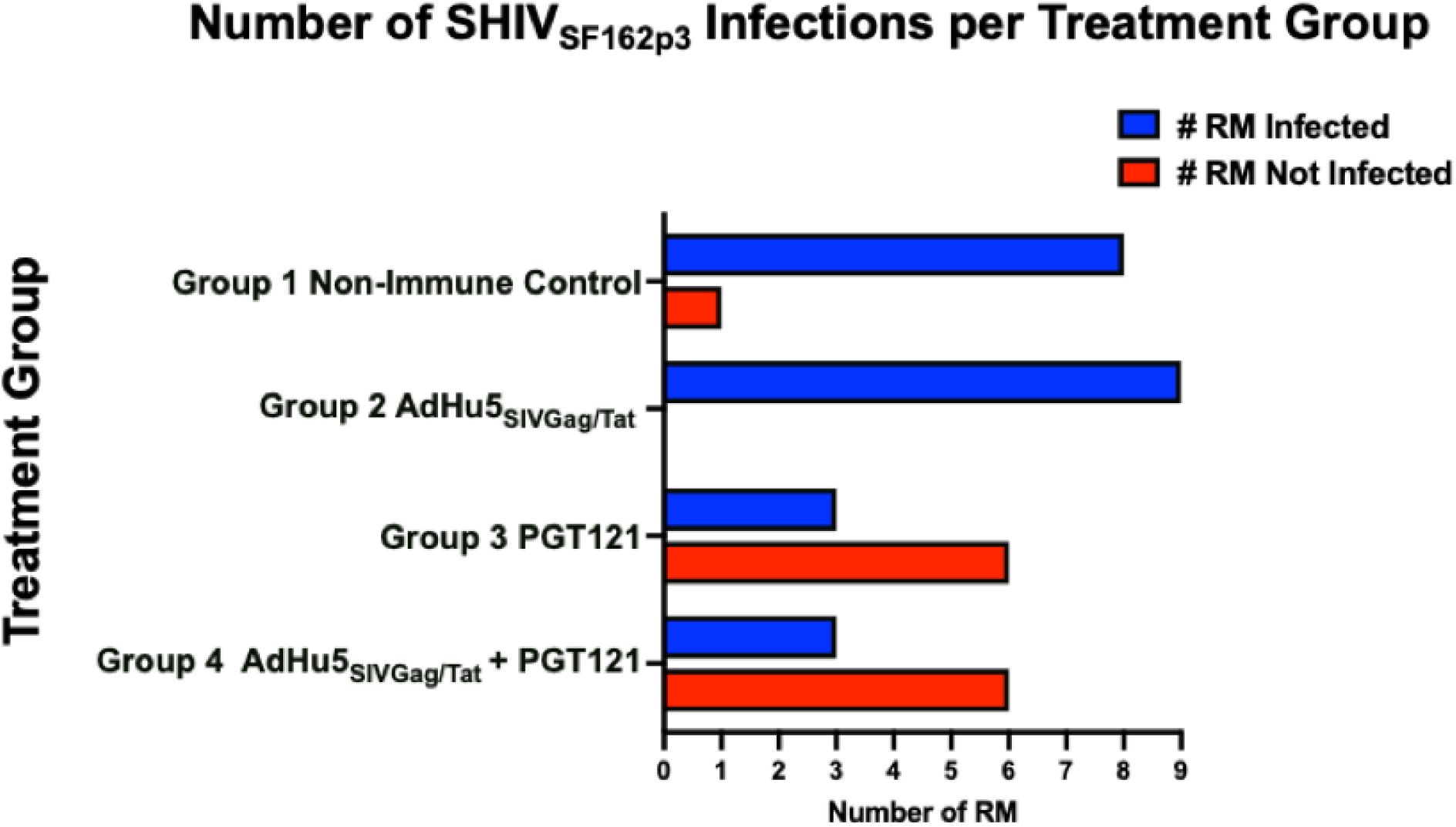
Effect of treatment interventions on the number of RM infected by SHIV_SF162p3._

**Table 1.**
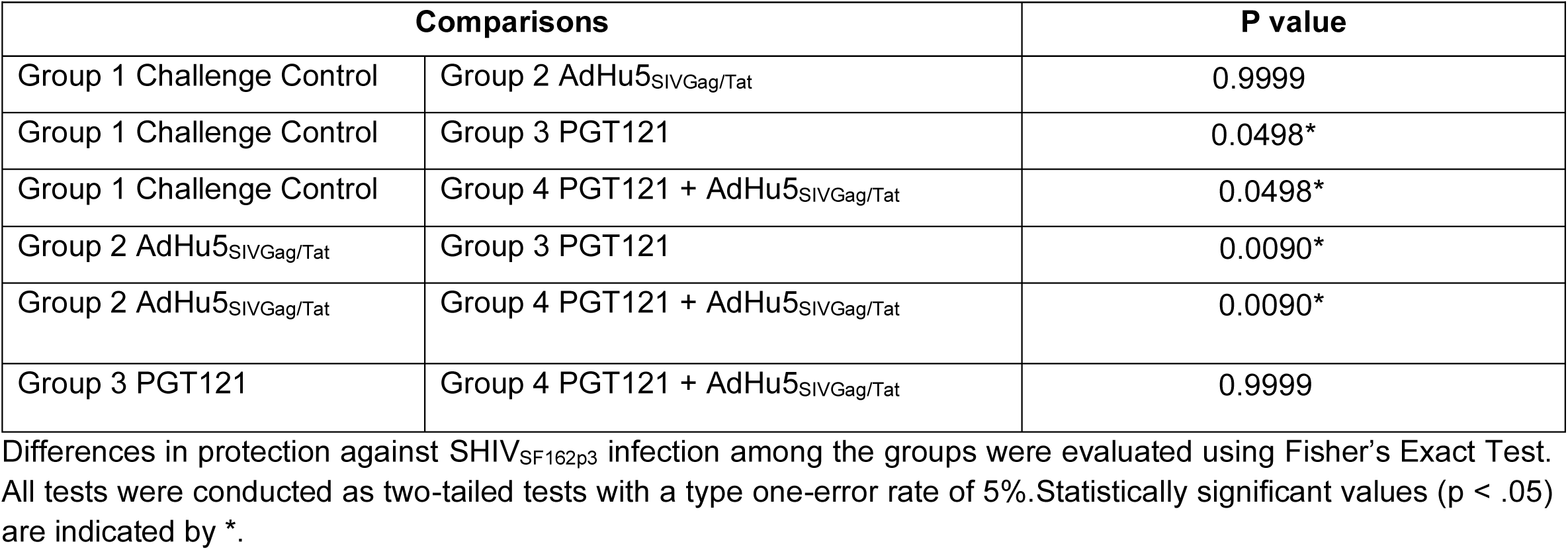
Differences in Protection against SHIV_SF162p3_ Infection.

### Constitutive rectal CD4+α4β7+Ki-67+ T cell Frequencies Correlate with Susceptibility to SHIV_SF162p3_ Infection

Rectal CD4+ T cell frequencies three weeks before the SHIV_SF162p3_ challenge were not significantly different among the groups, regardless of immunization status (Figure S4). Surprisingly, subgroup analyses (Figure 3) showed that the six infected RM in Groups 3 and 4 that received PGT121 collectively had higher constitutive frequencies of rectal CD4+α4β7+Ki-67+ T cells three weeks before the SHIV_SF162p3_ challenge than RM completely protected against SHIV_SF162p3_ infection. This result was independent of AdHu5_SIVGag/Tat_ vaccination, as it occurred in Group 3, which was not vaccinated. Further, the one RM that remained uninfected in the control group (Group 1) had a statistically significant lower frequency of rectal CD4+α4β7+Ki-67+ T cells than the infected RM. Thus, infection versus protection at a limiting bnAb dose of PGT121 appears to be determined by the frequencies of constitutive, vaccination-independent, CD4+α4β7+Ki-67+ T cells in the rectal mucosa before challenge. This finding is consistent with earlier results reported by our team members in the SIV system [56] and with another SIV vaccine study in which CD4+α4β7+Ki-67+ T cells attenuated protection [63]. A corollary of this result is that the limiting dose of PGT121 protects CD4+α4β7+Ki-67+ T cells from infection in the six RM in Groups 2 and 4 that resisted the high-dose rectal challenge with SHIV_SF162p3_. This is a new finding that deserves further consideration in the design of bnAb passive immunization studies in RM.

**Figure 3.**
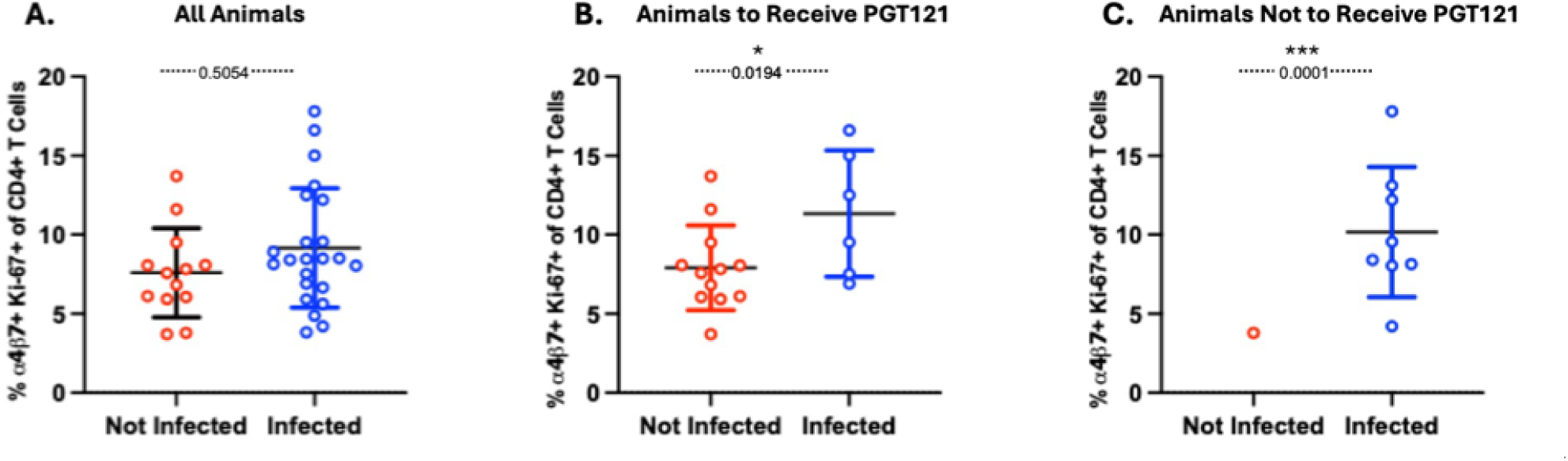
Frequencies of a4b7+Ki-67+ of CD4+ T-cells in rectal mucosa three weeks prior to challenge. The p values for panels A and B were determined by two-tailed t-test. The p value for the panel C was determined using one sample t-test using GraphPad Prism subroutines. Horizontal bars depict the mean ± SD.

### Three Distinct Patterns of Post-infection Control of Viremia are Elicited by bnAb, T cell Vaccine, or Combinatorial bnAb/T Cell Vaccine Interventions Before SHIV_SF162p3_ Challenge

The results shown in Figure 4 revealed three distinct patterns of post-infection control of viremia, including setpoint viremia. The first pattern is shown in Figure 4A, where RM immunized three times with AdHu5 _SIVGag/Tat_ exhibited classical post-infection control of viremia by T cell vaccination. This pattern of post-infection virus control is characterized by decreased peak viremia (Table 2) and decreased viral loads after challenge (Table 3). The slope of the setpoint viral load curve (weeks 5 to 14) was not significantly different from zero (P=.128 by linear regression, GraphPad Prism), indicating stable post-infection control of viremia. This pattern of vaccine-elicited post-infection control of viremia is a general characteristic of “T cell” vaccination strategies, in which there is no protection against acquisition but enhanced control of viremia post-infection [64–70].

**Figure 4.**
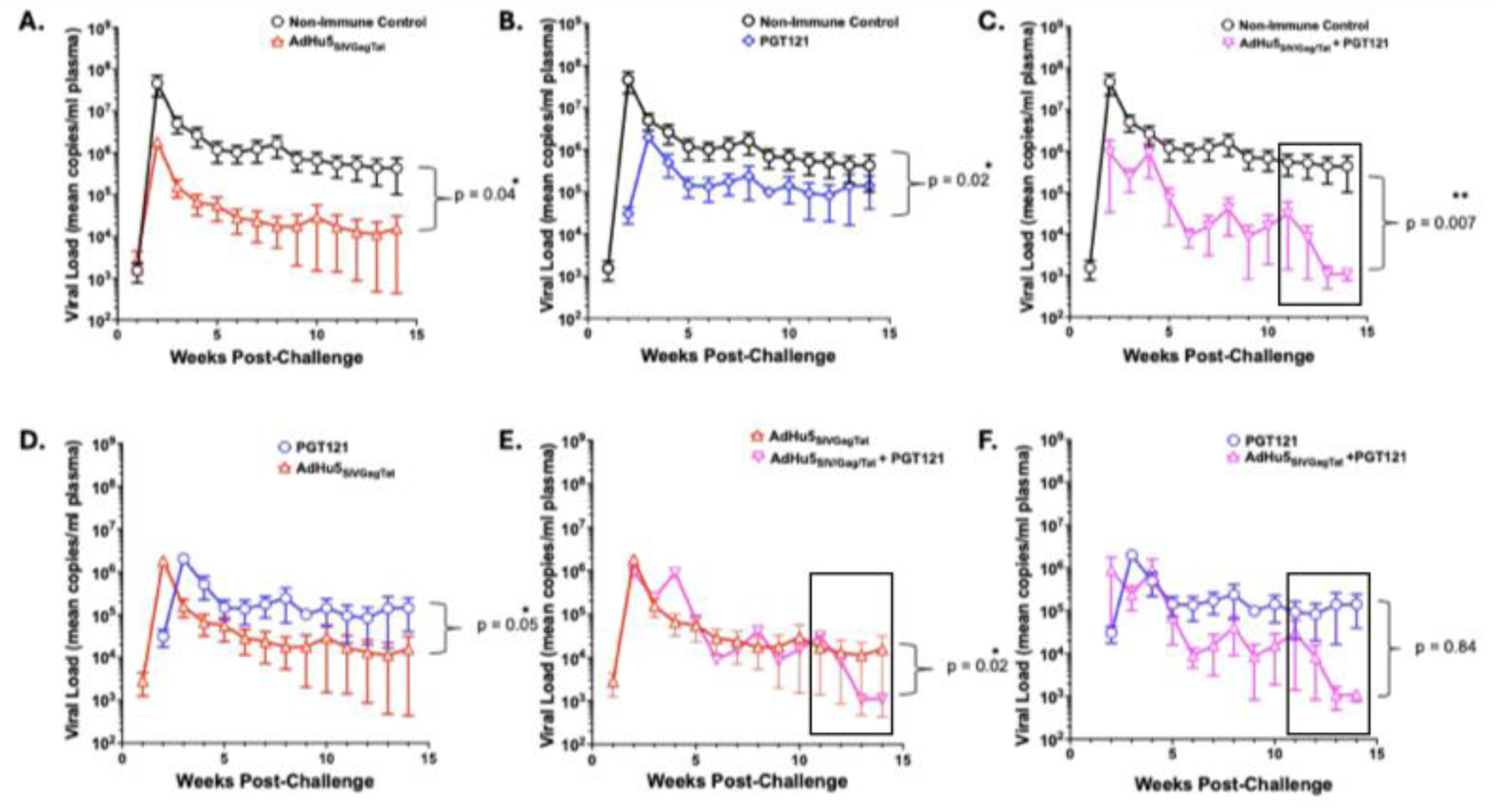
Distinct patterns or post-infection viral control elicited by immunization with AdHu5_SIVGag/Tat_, passive immunization with bnAb PGT-121, or with both treatments. P values are shown for differences in area under the curves for viral loads determined in Table 3. The boxes depict the weeks 11-14 inflection points for the group receiving AdHu5_SIVGag/Tat_ + PGT121 vs the groups receiving either intervention alone.

**Table 2.** Differences in Peak Viremia.

| Comparisons |  | P value |
|---|---|---|
| Group 1 Challenge Control | Group 2 AdHu5<sub>SIVGag/Tat</sub> | 0.0002* |
| Group 1 Challenge Control | Group 3 PGT121 | 0.0143* |
| Group 1 Challenge Control | Group 4 PGT121 + AdHu5<sub>SIVGag/Tat</sub> | 0.0009* |
| Group 2 AdHu5<sub>SIVGag/Tat</sub> | Group 3 PGT121 | 0.5203 |
| Group 2 AdHu5<sub>SIVGag/Tat</sub> | Group 4 PGT121 + AdHu5<sub>SIVGag/Tat</sub> | 0.0202* |
| Group 3 PGT121 | Group 4 PGT121 + AdHu5<sub>SIVGag/Tat</sub> | 0.1303 |

Differences in peak viremia among the groups were determined for log transformed data using two tailed ANOVA and pairwise t tests. One way ANOVA indicated differences among the groups (p = .0002). All tests were conducted as two-tailed tests with a type one-error rate of 5%. Statistically significant values (p < .05) are indicated by \*.

**Table 3.** Differences in Viral Load.

| **Comparisons** |  | **P value** |
|---|---|---|
| Group 1 Challenge Control | Group 2 AdHu5<sub>SIVGag/Tat</sub> | 0.04* |
| Group 1 Challenge Control | Group 3 PGT121 | 0.02* |
| Group 1 Challenge Control | Group 4 PGT121 + AdHu5<sub>SIVGag/Tat</sub> | 0.007* |
| Group 2 AdHu5<sub>SIVGag/Tat</sub> | Group 3 PGT121 | 0.05* |
| Group 2 AdHu5<sub>SIVGag/Tat</sub> | Group 4 PGT121 + AdHu5<sub>SIVGag/Tat</sub> | 0.02* |
| Group 3 PGT121 | Group 4 PGT121 + AdHu5<sub>SIVGag/Tat</sub> | 0.84 |

Areas under the curve (AUC) were determined using the viral loads for each experimental group using the method described in [1]. Kruskal-Wallis analysis indicated that the AUC estimate was different by vaccine group (p = .001), followed by six pairwise comparisons using the Wilcoxon Rank Sum Test shown in Table 1. All tests were conducted as two-tailed tests with a type one-error rate of 5%. Statistically significant values (p < .05) are indicated by \*.

The second pattern of post-infection viral control is shown in Figure 4B for the three RM infected after passive immunization with the partially protective dose of bnAb PGT121. This pattern features a significantly prolonged eclipse phase (days to first detectable viremia, Table 4), lower peak viremia (Table 2), and reduced viral load after challenge compared with the challenge control (Table 3). Notably, although passive immunization with PGT121 resulted in lower post-challenge viral loads than the challenge control group (Figure 4C, Table 3), these levels were still higher than in the RM immunized with AdHu5_SIVGag/Tat_ (Figure 4D, Table 3). The slope of the setpoint viral load curve (weeks 5 to 14) for the 3 RM infected after PGT121 passive immunization was not significantly different from zero (P=.562, linear regression, GraphPad Prism), indicating stable post-infection control of viremia. This pattern of post-infection control of viremia has been repeatedly observed in passive immunization studies using less potent neutralizing antibodies, such as the CD4-binding site-specific mAb b12, as well as in breakthrough infections following passive immunization with multiple bnAbs, including PGT121 [71].

**Table 4.** Differences in Eclipse Phase (First day of Viremia)

| Comparisons |  | P value |
|---|---|---|
| Group 1 Challenge Control | Group 2 AdHu5<sub>SIVGag/Tat</sub> | 0.9999 |
| Group 1 Challenge Control | Group 3 PGT121 | 0.0182* |
| Group 1 Challenge Control | Group 4 PGT121 + AdHu5<sub>SIVGag/Tat</sub> | 0.0182* |
| Group 2 AdHu5<sub>SIVGag/Tat</sub> | Group 3 PGT121 | 0.0045* |
| Group 2 AdHu5<sub>SIVGag/Tat</sub> | Group 4 PGT121 + AdHu5<sub>SIVGag/Tat</sub> | 0.0045* |
| Group 3 PGT121 | Group 4 PGT121 + AdHu5<sub>SIVGag/Tat</sub> | 0.9999 |

Differences eclipse phase viremia were dichotomized and evaluated using Fisher’s Exact Test. All tests were conducted as two-tailed tests with a type one-error rate of 5%.Statistically significant values (p < .05) are indicated by \*.

A third, unexpected pattern of post-infection control of viremia is shown in Figure 4 C for the three RM that became infected after immunization with AdHu 5 _SIVGag/Tat_ and passively immunized with PGT121, using the schedule shown in Figure 1. Immunization with AdHu 5 _SIVGag/Tat_ coupled with PGT121 passive immunization resulted in an increased eclipse phase (Table 4), decreased peak viremia (Table 2), and lower viral loads than the challenge control (Table 3). Surprisingly, there was a trend toward decreasing viremia in the viral setpoint window (weeks 5 to 14) for this group, as indicated by nonlinear regression analysis showing a slope distinct from 0 (P = 0.076, GraphPad Prism). There was a unique inflection point at week 11, where the slope of the week 11 to 14 viral load curve became significantly different from 0 by linear regression analysis (P = 0.026, GraphPad Prism). By contrast, the slopes of the 11- to 14-week viral load curves for the other intervention groups were not significantly different from 0 by linear regression analysis (P =0.840 for Group 2, P = 0.675 for Group 3, GraphPad Prism). Preliminary forecasting by autocorrelation analysis (JMP statistical software, SAS) suggests the viral load curve is trending toward fewer than 100 copies/ml shortly after the last observation week, although the data were too sparse to draw a firm conclusion without repeating the study using a longer observation window, as discussed below. Interestingly, the viral load curves were significantly reduced for RM receiving AdHu 5 _SIVGag/Tat_ + PGT121 vs those receiving AdHu 5 _SIVGag/Tat_ alone (Figure 4 E and Table 3, Group 1 vs Group 3). By contrast, the viral load curves were not statistically different for RM receiving AdHu5 _SIVGag/Tat_ + PGT121 vs those receiving PGT121 alone (Figure 4 F and Table 3, Group 2 vs Group 3), although there were differences in the inflection point observed in RM receiving AdHu5_SIVGag/Tat_ + PGT121. Taken together, these results are most consistent with three distinct patterns of post-infection viral control for each of the intervention groups, both alone and in combination. Possible mechanisms for these differences are discussed below.

## Discussion

This study investigated interactions between antibody-mediated protection against SHIV_SF162p3_ infection and vaccine-elicited T cell responses. The study was designed initially to evaluate the ability of vaccine-elicited CD4+ T cells to attenuate antibody-mediated protection against SHIV/SIV acquisition. This design was based on our previous vaccine studies [43, 53, 56] and a wider body of vaccine literature in NHPs [48, 52, 54, 55, 72] and humans [5, 44–46, 51], suggesting that vaccine-elicited CD4+ T cells are a double-edged sword in terms of protection against virus acquisition. On the one hand, vaccine-elicited CD4+ T cells are essential partners that provide necessary “help” to both B cells and CD8+ T cells in the adaptive immune response. On the other hand, subsets of vaccine-elicited CD4+ T cells are targets for HIV-1 infection. For an HIV-1 vaccine to work, it should elicit sufficient CD4+ T cell help to enable protective antibody and CD8+ T cell responses without compromising vaccine efficacy by creating excess CD4+ T cell targets for HIV-1 replication. We designed the study shown in Figure 1 to test this hypothesis. Because no current HIV-1 vaccines elicit bnAbs, bnAb-mediated protection was modeled by passive immunization with a limiting dose of bnAb PGT121 that partially protects against infection. A partially protective dose of PGT121 provided the most sensitive window to determine whether T cell vaccination affected protection against SHIV_SF162p3_ infection. A similar passive immunization strategy was used for PGT121 [73] to resolve whether Fc-mediated effector function contributes to its protection against a high-dose vaginal challenge with SHIV_SF162p3_ [73, 74]. T cell vaccination was modeled using AdHu5_SIVGag/Tat_, which is one of the vaccines used in our previous study showing that activated CD4+CCR5+ T cells in the rectum predict increased SIV acquisition in RM [56]. We used PGT121 as the bnAb and a single, rectal high-dose SHIV_SF162p3_ challenge as no SIV-specific bnAbs existed when the study was designed. Unlike our previous study [56], a single rectal high-dose SHIV_SF162p3_ challenge was used to simplify bnAb dosing. Three significant results emerged from this study. First, AdHu5_SIVGag/Tat_ vaccination did not affect PGT121-mediated protection against SHIV_SF162p3_ infection. Second, vaccine-independent frequencies of rectal CD4+α4b7+Ki67+ T cells measured three weeks before the SHIV_SF162p3_ challenge correlated with susceptibility to infection. Third, combinatorial bnAb/T-cell vaccination using PGT121 plus AdHu5_SIVGag/Tat_ revealed distinct patterns of post-infection control of viremia, suggestive of synergistic interactions between bnAbs and T-cell vaccination in controlling viral viremia. Several aspects of each finding deserve further discussion.

Within the parameters of the study, AdHu5_SIVGag/Tat_ T-cell vaccination neither enhanced nor attenuated the ability of a limiting dose of PGT121 to block SHIV_SF162p3_ infection. Regardless of PGT121 administration, AdHu5_SIVGag/Tat_ T-cell vaccination Gag-specific CD4+ and CD8+ T-cell responses in PBMC and rectal biopsies at levels enabling post-infection control of viremia, as seen in our previous study [56]. This finding does not rule out the possibility that other T-cell vaccination strategies might elicit CD4+ T-cell responses that attenuate antibody-mediated protection or dampen post-infection control of viremia. Several studies [49, 52–55, 72], including ours [53] have correlated vaccine-elicited CD4+ T cells with attenuation of protection associated with non-neutralizing antibody (nnAb) responses, which only reduce acquisition in repeat, low-dose SHIV/SIV challenges [52–54, 72]. As noted above, passive immunization studies using non-neutralizing mAbs have failed to prevent acquisition following high-dose SHIV_SF162p3_ challenges [75, 76], in stark contrast to many studies showing that bnAbs consistently block high-dose SHIV/SIV challenges. The data presented above extends this view to monoclonal bnAb doses that only partially protect against SHIV_SF162p3_ acquisition. Although T cell immunization with AdHu5_SIVGag/Tat_ did not increase SHIV_SF162p3_ acquisition or attenuate protection against virus acquisition, these studies shed new light on CD4+ T cell subsets that favor infection.

The increased HIV-1 acquisition observed in the STEP/Phambili efficacy trials [5, 44, 45] spurred further interest in defining the populations of vaccine-elicited CD4+ T cells associated with increased infection and decreased vaccine efficacy. A series of studies in human volunteers in HIV-1 vaccine trials [48, 77, 78] showed that AdHu5 vector-specific CD4+ T cells are more susceptible to HIV-1 infection than ALVAC vector-specific CD4+ T cells. Similarly, AdHu5-specific CD4+ T cells arising from natural adenovirus infection are also very susceptible to HIV-1 infection in vitro compared to CMV-specific CD4+ T cells [77, 79]. A study comparing the relative susceptibilities of other pathogen-specific CD4+ T cells to HIV-1 infection in vitro established the following rank order of susceptibility (more to less susceptible): AdHu5*, C. Albicans*, >> tetanus toxoid, HSV-2, >>> CMV [78, 79]. Subset and transcriptomic analyses suggest that the relative susceptibilities of CD4+ T cells specific for AdHu5, *C. Albicans*, tetanus toxoid, and HSV-2 to HIV-1 infection is associated with a robust CD4+ Th17 profile compared with the marked resistance of CMV-specific CD4+ T cells to HIV-1 infection that lack a Th17 profile [48, 77–79]. Notably, the relative resistance of CMV-specific CD4+ T cells to HIV-1 infection was also associated with the secretion of anti-HIV-1 b-chemokines, CCL3 (MIP-1α), CCL4 (MIP-1β), and CCL5 (RANTES) [77–79]. Thus, there is a clear difference among antigen-specific CD4+ T cell subsets for their relative susceptibility to HIV-1 infection. A recent study in RM associated strong Th-1 biased mucosal vaccine-specific CD4+ T cells with diminished vaccine efficacy [54]. Consistent with the HIV-1 infection susceptibilities of human antigen-specific CD4+ T cells, the vaccine elicited CD4+ T cells associated with reduced vaccine efficacy in RM secreted IFN-g and were strongly CCR5+ and did not secrete the antiviral b-chemokine, MIP-1β [54]. Vaccine-elicited protection was abrogated when frequencies of antigen-specific Th1-biased CD4+ T cells in the blood were above approximately 1.4%. This result is consistent with other vaccine studies in RM using repeat, low-dose mucosal SIV [52, 53] or SHIV [53, 54] challenges where circulating antigen-specific CD4+ IFN-γ+ T cells above a threshold of approximately 0.5% to 1.5% were associated with decreased protection against infection at lower levels of anti-Env antibodies. Our study further supports an antigen-specific CD4+IFN-γ+ T cell “threshold’ model for increased HIV-1/SHIV/SIV infection.

As shown in Figure 2, T cell vaccination with AdHu5_SIVGag/Tat_ did not affect protection against SHIV_SF162p3_ infection in the presence or absence of PGT121. This outcome is consistent with the “threshold” studies [52–54] in that the frequencies of circulating antigen-specific CD4+IFN-γ+ T cells were never above 0.35%, which is below the threshold predicted from the prior NHP studies [52–54] (Figure S1, panel B). The low frequencies of circulating vaccine-specific CD4+ IL-17A+ T cells are also consistent with the inability of AdHu5_SIVGag/Tat_ vaccination to affect protection against SHIV_SF162p3_ acquisition by a low dose of PGT121. These results suggest that the vector and immunogen choice should be optimized to induce vaccine-specific CD4+ T cells with an HIV-1 “resistant” phenotype while still providing T cell help. In this regard, it is interesting that vaccine-elicited CD4+ T follicular helper cells (Tfh) were not associated with diminished vaccine efficacy [54]. As suggested in [54], a vaccine that selectively favors antigen-specific CD4+ Tfh over Th-1 biased CD4+ T cells deserves further pursuit to minimize CD4+ T cell-mediated attenuation of protection.

While AdHu5_SIVGag/Tat_ vaccination did not affect the ability of PGT121 to protect against a high-dose SHIV_SF162p3_ challenge, an unexpected observation suggested that constitutive rectal CD4+α4β7+Ki-67+ T cells play a significant role in the susceptibility of RM to SHIV_SF162p3_ infection, regardless of vaccination with AdHu5_SIVGag/Tat_. While there were no overall differences in rectal CD4+α4β7+Ki-67+ T cells among infected vs. non-infected RM three weeks before the challenge (Figure 3, panel A), a direct association between rectal CD4+α4β7+Ki-67+ T cell frequencies and SHIV_SF162p3_ infection was found in RM from Groups 3 and 4 that received PGT121 (Figure 3, panel B). Infected RM had a higher frequency of constitutive CD4+α4β7+Ki-67+ T cells in the rectum three weeks before the SHIV_SF162p3_ challenge than RM in these groups protected against infection. Thus, constitutive CD4+α4β7+Ki-67+ T cells appear to attenuate a limiting dose of PGT121 to protect against infection. However, post-infection control of viremia was observed in Groups 3 and 4 RM that became infected despite receiving PGT121. This conclusion is supported by the finding that one RM in the control group (Group 1) was not infected by the high-dose rectal SHIV_SF162p3_ challenge.

Interestingly, the one uninfected in the SHIV_SF162p3_ challenge control group (Group 1) had a frequency of rectal CD4+α4β7+Ki-67+ T cells statistically lower than those of rectal CD4+α4β7+Ki-67+ T cells among the RM in Group 1 determined by a one-sample t-test. These results are consistent with the “threshold” hypothesis suggested above. The results are also consistent with our earlier repeat, low-dose SIV_mac239_ challenge study, where the frequencies activated rectal CD4+ T cells were associated with increased early viremia [56]. It should be noted that we did not observe increased early viremia in the current study, but this could be due to differences between the challenge strategy (high-dose vs. repeat, low-dose), the virus challenge stocks (SIV_mac239_ vs SHIV_SF162p3_), or both. Regardless of these differences, our current study further implicates activated CD4+ T cells at the site of virus entry as a determinant of infection susceptibility. We suggest that future RM studies should evaluate the frequencies of activated CD4+ T cells and susceptibility to SIV/SHIV infection, particularly in the context of antibody levels that do not completely prevent acquisition.

These observations augur for identifying immunogens that provide the CD4+ T cell help required for antibody and CD8+ T cell responses resistant to HIV-1/SIV/SHIV infections. Choosing which CD4+ T cell phenotype to target for these properties is made more difficult by the phenotypic heterogeneity of CD4+ T cells that become infected very early after virus exposure. This problem is evident in the studies suggesting that both Th1-biased CD4+ T cells [53, 54] or Th17 CD4+ T cells [48, 77–80] are associated with increased HIV-1 or SIV/SHIV infections, depending on the study. A very recent study in the RV254/SEARCH 010 acute infection cohort identified multiple CD4+ T cell phenotypes that become infected in the earliest stages of HIV-1 infection (Fiebig stages I-V) that share the proliferation and memory signatures as well as high levels of CCR5 [81, 82], which is consistent with the current study and a substantial literature [48, 50, 53, 54, 77–80]. Interestingly, the precise phenotypes of these early HIV-1 infected CD4+ T cells varied with the Fiebig stage with a preponderance of proliferating CD4+ T cells in Fiebig Stages I and II [81]. Given the rapidly expanding genomic and flow cytometry tools to characterize CD4+ T cell subsets, it is likely that future studies will identify a vaccination strategy that selectively elicits a CD4+ T cell subset that provides T cell help without becoming infected with HIV-1. In this regard, it is interesting to note that in humans, CMV-specific CD4+ T cells are very resistant to HIV-1 infection [77–79] and that a rhesus macaque cytomegalovirus vectored SIV T cell vaccine (rhCMV-SIV) is the only example of vaccination to successfully eliminate an early SIV infection associated with vaccine-specific CD8+ T cells [21–25, 42]. These findings warrant additional studies to design HIV-1 T cell vaccination strategies to elicit antigen-specific CD4+ T cells that are relatively resistant to infection by HIV-1.

Potentially, the most important finding of this study is the unique pattern of post-infection control of viremia elicited by the PGT121/AdHu5_SIVGag/Tat_ combinatorial bnAb/T cell immunization strategy (Figure 4, panels C and D). T cell vaccination with AdHu5_SIVGag/Tat_ or passive immunization with a limiting dose of PGT121 recapitulated patterns of post-infection control reported previously in NHP for T cell vaccination [56, 64–69] and bnAb passive immunization [83–85], respectively. By contrast, the pattern of post-infection control of viremia elicited by combinatorial PGT121/AdHu5_SIVGag/Tat_ immunization was distinct from either intervention alone. There was an increased time to first viremia, a delayed and reduced peak viremia, and a potent steady-state viremia that began to decay over the last four weeks of the study. These results suggest additive and potentially synergistic interactions between bnAbs and T cell vaccination in post-infection control of viremia. The increased time to first viremia is similar to that observed for passive immunization with PGT121 alone. The robust post-infection control of viremia is similar to that observed for immunization with AdHu5_SIVGag/Tat_ alone. In this sense, the effects of each single intervention are approximately additive in response to combinatorial PGT121/AdHu5_SIVGag/Tat_ immunization. The unique decay of steady-state viremia from weeks eleven to fourteen in the PGT121/AdHu5_SIVGag/Tat_ immunization group suggests combinatorial synergy not seen with either immunization alone. This observation is consistent with a recent study demonstrating synergy between bnAbs and antigen-specific CD8+ T cells in controlling HIV-1 replication in vitro [19].

Our study extends two previous studies in rhesus macaques [20, 86] that examined the relationship between bnAb prophylaxis and T-cell vaccination for protection against SHIV/SIV. Although the three studies differ in many respects, several common themes emerged, supporting continued development of bnAb/T-cell vaccination strategies. All three studies used limiting bnAb doses that partially protected against SHIV/SHIV infection when administered alone, providing optimal windows to evaluate the effect of T-cell vaccination on bnAb-mediated protection. Prior studies suggest that sub-protective levels of bnAbs at the time of virus exposure reduce the size of the viral inoculum, delay the eclipse phase, and modestly decrease viral setpoints [87], providing fertile ground for increased post-infection control of viremia by T-cell vaccination. Across the three studies, T-cell vaccination had no apparent effect, positive or negative, on protection against viral acquisition. As expected, T-cell vaccination alone did not prevent infection in any of the studies. However, in combination with bnAb prophylaxis, T-cell vaccination blunted peak viremia in the earliest study [86], enhanced viral replication arrest in the second study [20], and accelerated viral setpoint decay to near-background levels in our study. Synergistic interactions between bnAbs and vaccine-elicited CD8+ T cells in replication arrest and accelerated viral decay are strongly suggested in [20] and in our study. These studies support a dual-modality HIV-1 vaccine strategy to blunt the pandemic, in which bnAbs protect against infection in most individuals and synergistic interactions between bnAbs and CD8+ T cells arrest or clear infections in those for whom the bnAb titers were too low to prevent infection. These results fit the emerging picture of early bnAb interventions after acute infection and in structured acute-treatment interruption studies describing vaccinal effects, where bnAbs and virus-specific CD8+ T cells can result in profound control of infection in some individuals without subsequent ART. In this vein, studies of murine retroviral infections suggest that antigen cross-presentation of virus-antibody immune complexes by dendritic cells enhances virus-specific CD8+ T cell responses that control viral replication. In other studies, we have developed a mathematical model based on this type of antigen cross-presentation that accurately models the accelerated viral decay reported above and is being used to further test the vaccinal hypothesis in our combinatorial bnAb T-cell vaccination system.

While our results are consistent with interactions between antibody-mediated vaccinal effects and deliberate T cell vaccination, they remain hypothetical given the limitations of our original study design. Our study was designed to determine whether T cell vaccination affected protection against SHIV_SF162p3_ infection mediated by a limiting dose of bnAb PGT121. Accordingly, we did not anticipate differences in post-infection control of viremia among Groups 2, 3, and 4. Thus, we did not plan for CD8+ T cell depletion in infected RM in Groups 2, 3, and 4 to determine whether this subset mediates post-infection control. Similarly, we designed our original study to follow infection for fourteen weeks post-challenge, which is sufficient for cryptic infections to emerge in the PGT121-treated groups. However, the fourteen-week endpoint limited our ability to determine whether the accelerated decay in post-infection viremia observed in Group 4 would reach background levels or remain stable over a much longer time, as reported for bnAb interventions leading to CD8+ T cell-mediated control of post-infection viremia [15, 16, 88]. Another limitation of our study is that we have not conducted extensive tissue analyses to identify transcriptomic signatures associated with the different experimental outcomes. Although we collected appropriate tissues for transcriptomic analyses, we elected to bank them for use in a larger multi-center study to test the hypothesis that bnAb-mediated vaccinal effects and T cell vaccination synergizes to mediate a unique pattern of robust post-infection control of viremia.

Despite the above limitations, our studies demonstrate previously unknown interactions among bnAbs, T-cell vaccination, and pre-existing bnAbs that influence post-infection control of viremia in the SHIV_SF162p3_ challenge model. These results support an HIV-1 vaccination strategy that elicits bnAbs that block infection by most circulating variants and also synergize with vaccine-elicited T cells to enhance post-infection control of viremia in individuals whose bnAb responses still interact with the virus without preventing acquisition. This approach could mitigate the consequences of infection for individuals infected with neutralization-sensitive HIV-1 variants that are neutralized only at titers above the transmission threshold, as reported in the recent AMP studies on HIV-1 prevention by bnAb VRC01 [9–11].

## Materials and Methods

### Animals

Thirty-six healthy, SIV-uninfected, Mamu-B*08- and B*17-adult Indian rhesus macaques (macaca mulatta) (RMs) were used in this study. Half of the study population were Mamu-A*01-positive which was evenly distributed through the treatment groups. All procedures were approved by the Emory University Institutional Animal Care and Use Committee (IACUC) and animal care facilities are accredited by the U.S. Department of Agriculture (USDA) and the Association for Assessment and Accreditation of Laboratory Animal Care (AAALAC) International.

### AdHu5_SIVGag/Tat_ vaccine

The adenovirus vector component of AdHu5_SIVGag/Tat_ was derived from the human serotype 5 (AdHu5). The E1- and E3-deleted AdHu5 vector expressed a Gag-Tat fusion of SIV_mac239_ (AdHu5_SIVGag/Tat)._ The AdHu5_SIVGag/Tat_ vaccine was generated, rescued, expanded, purified, titrated and quality controlled as described [91]. Its immunogenicity was described in the previous RM study by our coauthors [56].

### Broadly neutralizing antibody PGT121

The PGT121 bnAb was produced at PlantVax Corporation (Rockville, MD) in tobacco plants by *Agrobacterium*-mediated transient transfection of *Nicotiana benthamiana* and purified as described in [92]. Multiple studies of *Nicotiana*-produced PGT121 by our coauthors have described its pharmacokinetics [61, 62] and shown it to be non-immunogenic in RM [61, 62] as well as consistently protective against infection with SHIV_SF162p3_ in RM [62, 93, 94]. The limiting dose of PGT121 (1.0 mg/kg) used to protect approximately sixty percent of RM from SHIV_SF162p3_ infection in this study was based on prior studies from the coauthors [61, 62, 93, 94]. PGT121 serum concentrations were determined by ELISA using plates coated with a conformationally constrained gp120 -CD4 fusion protein (FLSC) [95] expressed in mammalian cells (HEK293). The conformationally constrained FLSC protein reacts strongly with PGT121 that recognizes a N332 glycan-V3 epitope [96, 97], which is stabilized by the intramolecular binding of gp120 to CD4 in FLSC. Serial dilutions of a PGT121 standard or sera from RM passively immunized with PGT121 were incubated and washed as described [53]. Specific binding was determined by incubating the plates with a rabbit anti-human IgG1 hinge monoclonal antibody (clone RM117, ThermoFisher, Inc.) followed by six additional washes with ELISA buffer, and detection using a goat anti-rabbit IgG alkaline phosphatase conjugate (ThermoFisher, Inc.). Neither the rabbit anti-human IgG1 hinge monoclonal antibody, nor the goat anti-rabbit IgG-alkaline phosphatase conjugate were reactive with RM IgG in preliminary studies. PGT121 half-lives were estimated using sera collected over weeks 37-40 of the study by non-linear curve fitting (one phase decay) using Prism (GraphPad Software, Boston, MA).

### Vaccination and antibody infusion protocol

Of the thirty-six animals assigned to the study, eighteen animals received three immunizations of AdHu5_SIVGag/Tat_ performed at day 0, 16 weeks, and 32 weeks as detailed in Figure 1. All used vectors expressed codon optimized sequences of SIV_mac239_. Immunizations comprised of the AdHu5 vector expressing SIVGag/Tat were injected intramuscularly (IM) at the dose of 10^11^ virus particles per RM. Nine animals from the vaccinated animals and nine of the untreated animals received and intravenous infusion of 1 mg/kg of the broadly neutralizing antibody PGT121. Nine animals remained untreated.

### Virus challenge

One month following the final immunization, all RMs were challenged with a single intrarectal dose of a 1:10 dilution of SHIV_SF162p3_ that was provided by Dr. Ranajit Pal, ABL Inc., and Dr. Nancy Miller, NIAID, NIH. The challenge stock, SHIV_SF162p3_ (R922 derived) Harvest 3 dated 9/12/2016, had the following specifications: 182.79 ng/ml p27, >10^9^ RNA copies/ml, and 2032 TCID50/ml (tittered in rhesus PBMC).

### Plasma viral load determination

Quantitative real-time reverse-transcriptase (RT)-PCR assay to determine SHIV_SF162p3_ viral load was performed as previously described [89, 98]. The sensitivity of the assay is 60 copies/ml of plasma.

### Tissue collection and processing

Peripheral blood mononuclear cells (PBMCs) were isolated by gradient centrifugation. Procedures for lymph node (LN) biopsy, rectal biopsies (RB), as well as isolation of lymphocytes from the obtained samples were performed as previously described [89, 90].

### Immunophenotyping and flow cytometry

Multi-color flow cytometric analysis was performed on mononuclear cells isolated from blood, and mucosal tissues (RB) according to standard procedures using human monoclonal antibodies that were found to cross - react with RMs. The cocktail of antibodies for surface staining included viability dye efluor780 from eBioscience; CD8 BUV495 (3B5), CD3 BUV395 (SP34-2), CCR5 APC (3A9) from BD Bioscience; HLA-DR PerCP-Cy5.5 (L243), PD-1 BV421 (EH12.2H7), CD4 BV785 (OKT-4), CD25 BV605 (BC96) from Biolegend; and anti-a4b7 PE (A4B7R1) from the Nonhuman Primate Reagent Resource. The cocktail for intracellular staining included Ki-67 Alexa488 (B56) from BD Bioscience and FoxP3 PEDazzle594 (206D) from Biolegend. Flow cytometric acquisition and analysis of samples was performed on at least 100,000 lymphocytes on a LSRII flow cytometer driven by the DiVa software package (from BD Biosciences, San Jose, CA). Analysis of the acquired data was performed using FlowJo software (Tree Star, Inc., Ashland, OR).

### Tetramer staining

MHC class I tetramers were prepared and conjugated to APC fluorophore by the NIH Tetramer Core Facility as previously described (Altman 1997). The level of SIV-specific CD8^+^ T cells was assessed using soluble tetrameric Mamu-A*01 MHC class I tetramers specific for SIV_mac239_ immunodominant epitopes Gag181-189 CM9 (CTPYDINQM) in the vaccinated, Mamu-A*01+ RMs. Lymphocytes isolated from blood were incubated with conjugated tetramer, along with surface antibody conjugates, and analyzed for tetramer and surface marker expression using an Fortessa Flow Cytometer (from BD Biosciences, San Jose, CA) equipped with FACS DiVa software.

### Multifunctional assessment of SIV-specific T cell responses

The function of SIV-specific CD8^+^ T cells was assessed by flow cytometry after stimulation with peptide pools of 15-mers (overlapping by 11 amino acids) spanning the SIV_mac239_ Gag proteins as described in (Betts, 2006; Cervasi 2013), Peptides were prepared from peptide stocks obtained from the NIH AIDS Research and Reference Reagent Program (Bethesda, MD), reconstituted in DMSO and pooled. All peptides were used at a final concentration of 2 µg/ml of each peptide. Staining for flow cytometry was described previously [56]. The unstimulated control was used as a background and the percentage of responding cells is depicted after background subtraction.

### Statistical Analyses and Tables

Additional statistical analyses are reported as indicated in the text or figure legends. Continuous or dichotomous data were evaluated as indicated using the appropriate subroutines in Prism (GraphPad, Boston, MA). All tests were conducted as two-tailed tests with a type one-error rate of 5%.

## Acknowledgments

The authors thank Ms. Robin Flinko, Ms. Qiang Lin, Ms. Meng Li, and Mr. Scott Foulke of the IHV for outstanding technical assistance. We also thank Mr. Kirk Easley of the Rollins School of Public Health, Emory University, for initial statistical support in analyzing viral load curves. We thank Dr. Ranajit Pal (ABL) and Dr. Nancy Miller (NIH) for the titrated virus challenge stock. We thank Dr. Yvonne Rosenberg (PlantVax) for the PGT121 bnAb. We thank Dr. Anthony L. DeVico of the Institute of Human Virology for project advice.

## Research Support

This research was supported by National Institutes of Health’s Office of the Director, Office of Research Infrastructure Programs, P51 OD011132 and U42 OD011023. This research was also supported by NIAID, NIH, P01AI162242; NIAID, NIH, 5R01AI187023; and grants from The Bill and Melinda Gates Foundation. Dr. Lewis was also supported by the Robert C. Gallo, M.D., Distinguished Professorship in Translational Medicine.

---

**Figure S1:**
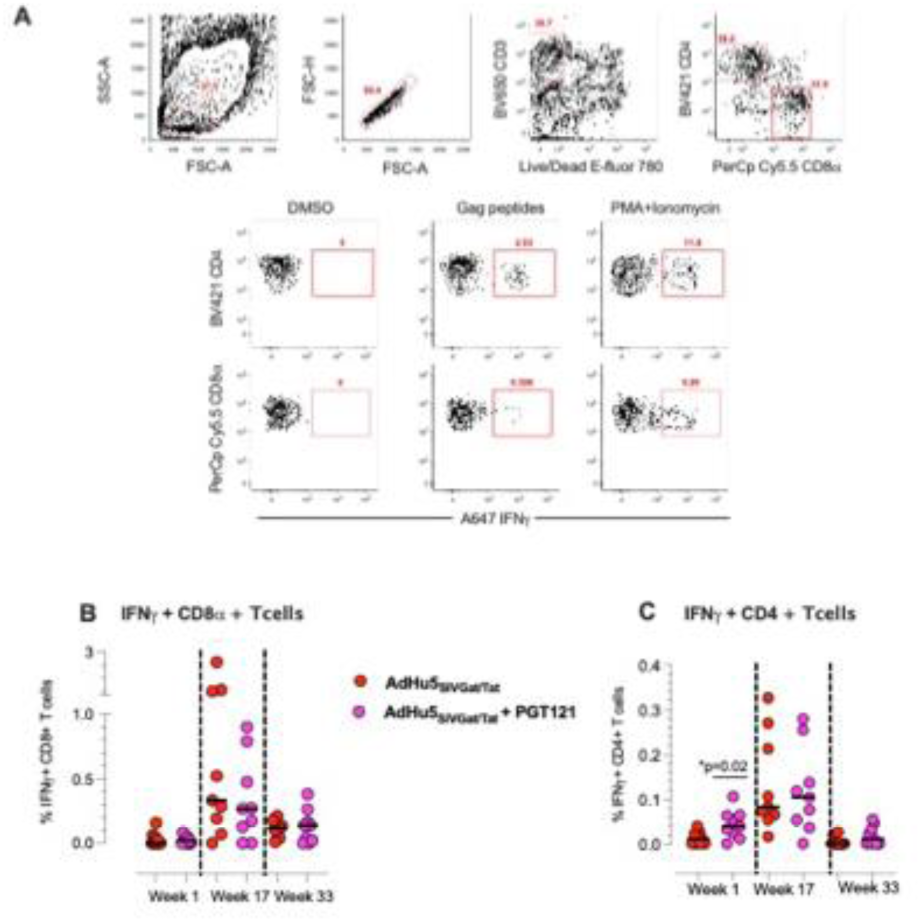
Quantification of SIV Gag specific T cells secreting IFN-γ post AdHu5_SIVGag/Tat_ immunizations using peptide pulse and intracellular cytokine staining. A) Representative flow cytometry plots show gating strategy used to identify IFN-γ secreting CD8+ and CD4+ T cell responses to the SIV gag peptide pool stimulations ex-vivo, as well as no peptide and PMA + Ionomycin stimulated negative and positive assay controls. Graphs summarize frequencies of B) IFN-γ + CD8+ T cells, C) IFN-γ + CD4+ T cells. Horizontal bars indicate median values. Two-tailed non-parametric Mann-Whitney U test was used to compare significance of difference between animals that received AdHu5_SIVGag/Tat_ immunizations alone or in those that received the PGT121 bNAb prior to SHIV_SF162p3_ challenge. * represents a p value <0.05.

**Figure S2:**
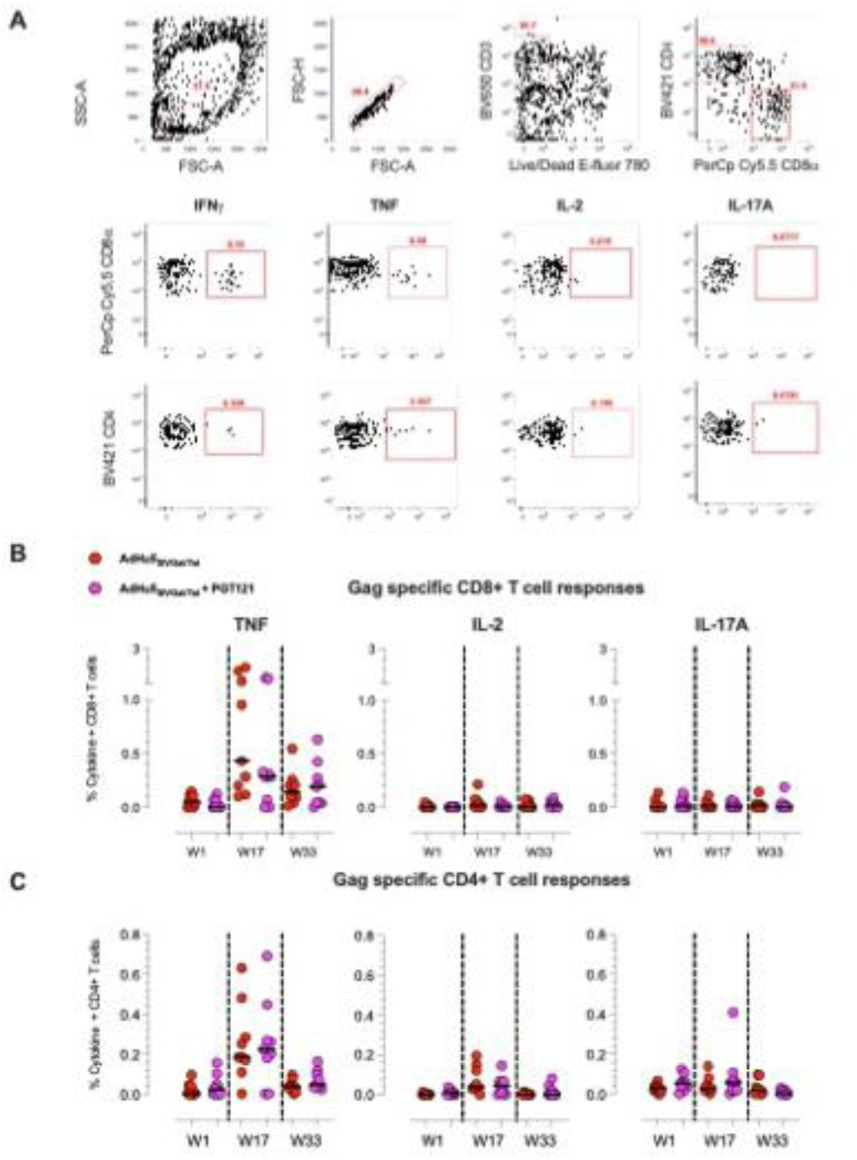
Quantification of SIV gag specific T cells secreting additional cytokines post AdHu5 vaccinations using peptide pulse and intracellular cytokine staining. A) Representative flow cytometry plots show gating strategy used to identify TNF, IL-2 or IL-17A secreting CD8+ and CD4+ T cell responses to the SIV gag peptide pool stimulations ex-vivo. Graphs summarize frequencies of B) TNF, IL-2 or IL-17A+ CD8+ T cells, and C) TNF, IL-2 or IL-17A+ CD4+ T cells. Horizontal bars indicate median values.

**Figure S3.**
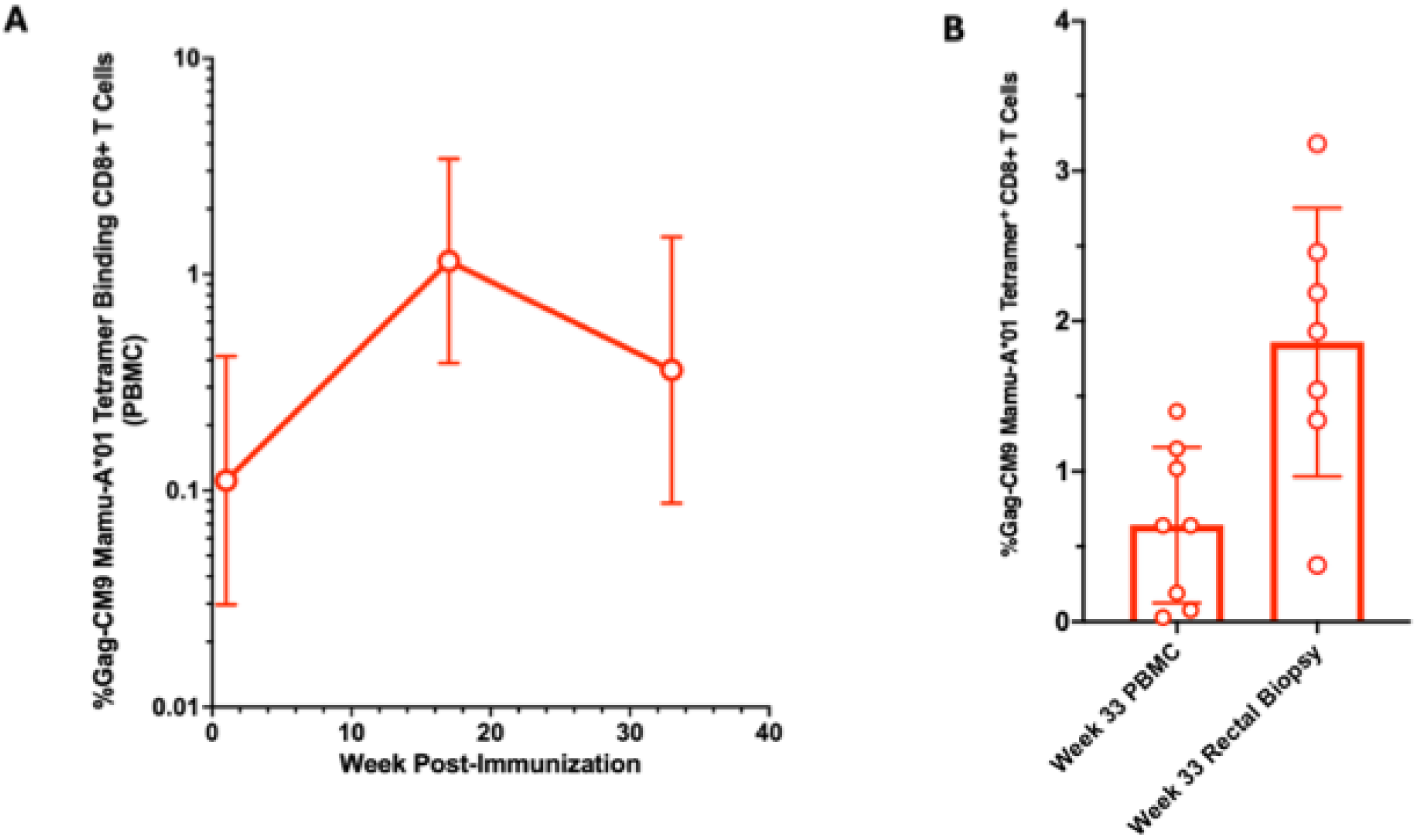
Tissue frequencies of %Gag-CM9 Mamu-A*01 Tetramer Binding CD8+ T Cells in RM vaccinated with AdHu5_SIVGat/Tat_. Panel A shows the frequencies of %Gag-CM9 Mamu-A*01 Tetramer Binding CD8+ T Cells in RM vaccinated with AdHu5_SIVGat/Tat_ in PBMC one-week post-vaccination. Panel B shows the week 33 frequencies of %Gag-CM9 Mamu-A*01 Tetramer Binding CD8+ T Cells in RM vaccinated with AdHu5_SIVGat/Tat_ PBMC and rectal biopsies.

**Figure S4.**
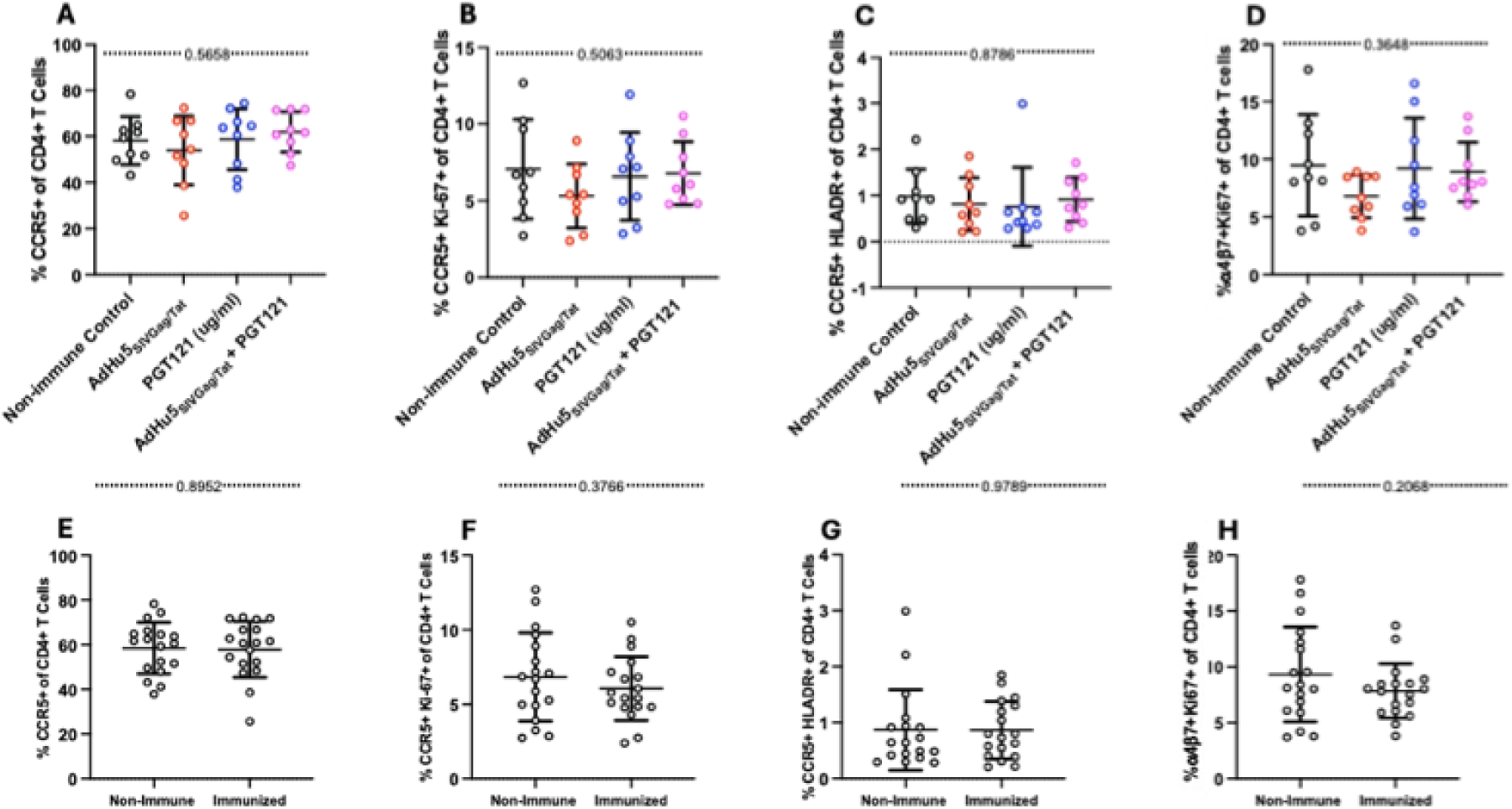
Frequency of activated and proliferating CD4+ T-cells in rectal mucosa three weeks prior to challenge. P values are shown between dashed lines for two tailed ordinary one-way ANOVA for the top four panels. P values determined by two-tailed t-test are shown for the bottom four panels. None of the differences were significant. Horizontal bars depict the mean ± SD.

**Figure S5.**
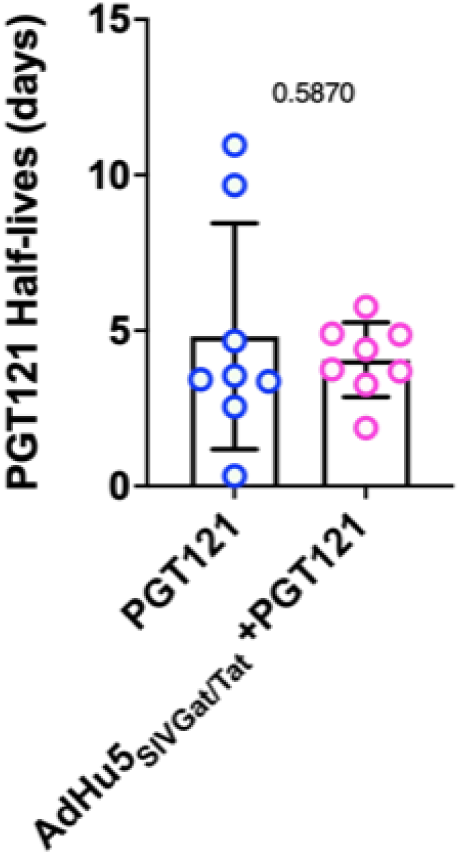
PGT121 serum half-lives for Group 3 (PGT121) and Group 4 (AdHu5_SIVGat/Tat_ + PGT121). Serum half-lives were calculated by non-linear regression (single exponential decay) using GraphPad Prism. The means and standard deviations are shown for each group. There was no significant difference between the groups determined by two tailed t-test, P=.5870. Due to sparse data, it was not possible to fit curves for one RM in each group and those data were excluded from the analysis.

## Notes

### Competing Interest Statement

The authors have declared no competing interest.

